# Doubling the resolution of fluorescence-lifetime single-molecule localization microscopy with image scanning microscopy

**DOI:** 10.1101/2023.08.23.554438

**Authors:** Niels Radmacher, Oleksii Nevskyi, José Ignacio Gallea, Jan Christoph Thiele, Ingo Gregor, Silvio O. Rizzoli, Jörg Enderlein

## Abstract

In this study, we integrate a single-photon detector array into a Confocal Laser Scanning Microscope (CLSM), enabling the combination of Fluorescence Life-time Single-Molecule Localization Microscopy (FL-SMLM) with Image Scanning Microscopy (ISM). This unique combination delivers a two-fold improvement in lateral localization accuracy for SMLM while maintaining its simplicity. Moreover, the addition of lifetime information from our CLSM eliminates chromatic aberration, particularly crucial for achieving few-nanometer resolution in SMLM. Our approach, named Fluorescence Lifetime ISM-SMLM (FL-iSMLM), is demonstrated through direct Stochastic Optical Reconstruction Microscopy (dSTORM) and DNA Point Accumulation for Imaging in Nanoscale Topography (DNA-PAINT) experiments on fluorescently labelled cells, showcasing both resolution enhancement and fluorescence lifetime multiplexing capabilities.

## 1 Introduction

Over the past decades, super-resolution microscopy has become a fundamental tool in bioimaging, finding manifold applications in biophysics and cell biology. One of the most popular and powerful super-resolution modalities is Single-Molecule Localization Microscopy (SMLM). This encompasses techniques like Photoactivated Localization Microscopy (PALM) [1], (direct) Stochastic Optical Reconstruction Microscopy (STORM [2] and dSTORM [3]), Point Accumulation for Imaging in Nanoscale Topography (PAINT) [4], and its especially potent variant DNA-PAINT [5]. Nowadays, SMLM can achieve lateral resolutions in the single digit nanometer range [6], two orders of magnitude better than the diffraction limit of classical optical microscopy.

Typically, SMLM is based on a wide-field microscope, enabling the detection and localization of single molecules over a large field of view. Recently, we developed a variant of SMLM based on a rapid Confocal Laser-Scanning Microscope (CLSM), which combines the super-resolution of SMLM with Fluorescence Lifetime Imaging (FLIM) [7, 8]. The additional fluorescence lifetime information not only allows for target multiplexing in imaging but also facilitates the combination of SMLM with Metal-Induced Energy Transfer (MIET) imaging [9–11]. MIET uses the modulation of fluorescence lifetime by a nearby metal layer to achieve nanometer resolution along the optical axis. By merging FL-SMLM with MIET, we achieved an isotropic single-molecule localization with a few nanometers localization accuracy along all three dimensions.

In this paper, we synergize CLSM-based FL-SMLM with another cutting-edge high-resolution technique known as Image Scanning Microscopy (ISM) [12, 13]. ISM employs a blend of focused laser excitation and wide-field imaging, accomplished by augmenting a CLSM with an array detector. This setup enables the capture of a miniature image of the illuminated sample at each scanning position of the CLSM. Through meticulous image reconstruction [14–16], ISM effectively doubles the lateral resolution and quadruples image contrast by reducing the effective point spread function by a factor of two.

The resolution enhancement accomplished by ISM is akin to that achieved by Structured Illumination Microscopy (SIM) [17], which also employs non-uniform structured illumination and wide-field detection to double the lateral resolution. In the realm of SMLM, ISM directly enhances the accuracy of single-molecule localizations, thus significantly elevating the ultimate SMLM resolution. From a perspective rooted in physical optics, this mirrors the twofold increase in localization accuracy achieved through patterned illumination techniques like SIMPLE [18, 19], SIMFLUX [20], as well as others [21, 22]. However, ISM offers a substantial reduction in experimental complexity. Instead of illuminating the sample with variously positioned and oriented excitation patterns, ISM merely requires the augmentation of a CLSM with a compact array detector and the acquisition of scan images of the sample in the standard CLSM operational mode.

Many published variants of ISM employ rapid cameras for recording the fluorescence signal, which lack the capability for performing FLIM [23, 24]. However, Tenne et al. pioneered the integration of several Single-Photon Avalanche Detector (SPAD) detectors and fast multichannel Time-Correlated Single Photon Counting (TCSPC) electronics, thus introducing fluorescence lifetime ISM [25, 26]. Recent developments in SPAD array technology have further extended the possibilities of combining ISM with FLIM [27–30].

In this study, we demonstrate that the integration of a SPAD array detector, along with fast TCSPC electronics, into our CLSM system enables us to achieve doubled resolution in FL-iSMLM. After expounding the theoretical foundations of iSMLM, we describe its experimental implementation and meticulously validate and calibrate it using fluorescent point-spread function beads. Subsequently, we showcase its capabilities by imaging the intricate microtubule lattice within fixed COS-7 cells. Additionally, we introduce two-target FL-iSMLM imaging of actin and tubulin. The achieved lateral resolution, approximately *∼*5 nm, underscores the significance of FL-iSMLM, as it completely eliminates chromatic aberration. Finally, we demonstrate a biologically significant application by simultaneously imaging Homer1 and Bassoon proteins within the synaptic cleft of neurons through the use of DNA-PAINT-based FL-iSMLM.

## 2 Results

### 2.1 Principle of Image Scanning SMLM

Since the advent of Confocal Laser Scanning Microscopy (CLSM), it has been recognized that the lateral resolution of such instruments could theoretically be doubled by significantly reducing the size of the confocal aperture. This reduction would make the aperture’s diameter much smaller than the Airy disk of a point emitter. However, this approach leads to a substantial reduction in light throughput, as the majority of light collected from the sample would be excluded by the pinhole. The fundamental concept of Image Scanning Microscopy (ISM) [12, 13] involves substituting the pinhole and single-point detector of a traditional CLSM with an array of small pixel detectors. These detectors are each significantly smaller than the Airy disk (see Fig. 2(a)). In this setup, although each pixel captures only a small amount of light, no light is generally lost due to the extended coverage of the entire detector array. Therefore, when scanning the sample with a focused laser spot, each pixel in the array detector captures its own scan image. This process ideally doubles the lateral resolution compared to conventional CLSM. Nevertheless, due to parallax effects, the scan image captured by one pixel is slightly offset from that captured by another. ISM employs a sophisticated algorithm to realign all these scan images into a unified frame, yielding a lossless, doubly resolved final image [14, 31]. Image Scanning Single-Molecule Localization Microscopy (iSMLM) combines ISM with Single-Molecule Localization Microscopy (SMLM). This technique first records scan images of well-separated fluorescent emitters using an array detector. Then, it reduces the size of the single-molecule images by approximately a factor of 2 using the ISM algorithm. Finally, it localizes the molecules within the ISM image. The reduction in the size of single-molecule images (”superconcentration of light”, see Ref. [32]) directly results in improved localization accuracy, thereby nearly doubling the resolution of the resultant super-resolved SMLM image.

For the sake of completeness, in what follows we briefly recapitulate the theoretical principles of ISM [12–14, 24, 31] and how it improves SMLM. The iSMLM raw data takes the form of a four-dimensional array of intensities. Two dimensions pertain to the position ***ρ*** of the scanning focus on the sample, and the other two dimensions correspond to the pixel position **s** on the array detector. For instance, when collecting data from a single molecule located at lateral position ***ρ*** = **0** and axial position *z* relative to the focal plane, the four-dimensional dataset *I*(***ρ***, **s***|z*) is expressed as:

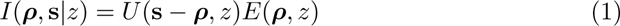

Here, *E*(***ρ****, z*) represents the three-dimensional excitation intensity distribution of the focused laser beam, and *U* (**s***−****ρ****, z*) is the three-dimensional Point Spread Function (PSF) for imaging on the array detector, analogous to the PSF in wide-field imaging. Note that the detection path is de-scanned, as depicted in Fig. 2(a), ensuring that the center of the array detector aligns with the center of the excitation focus. This explains the argument **s** *−* ***ρ*** in the function *U* .

In the following discussion, we assume that the imaged molecules reside in the focal plane, i.e., *z* = 0. The majority of the subsequent arguments remain applicable to molecules situated outside the focal plane, in which case the functions below must be evaluated at the specific axial position *z*.

To elucidate the structure of *I*(***ρ***, **s**), consider Fig. 1, which displays cross-sections of the excitation intensity distribution *E*(***ρ***) (central blue curve) and the detection efficiency *U* (**s** *−* ***ρ***) for a pixel at position **s** (orange curve). These illustrations assume diffraction-limited and aberration-free focusing and imaging. In this context, we employ the scalar approximation [*J*_1_(*q*_max_*ρ*)*/q*_max_*ρ*]^2^ for both excitation intensity and detection efficiency distributions. This function has been shown to be an excellent approximation of the actual PSF even for large values of numerical aperture, see e.g. Fig. 20 in Ref. [33]. Here, *J*_1_ represents the Bessel function of the first kind, and *q*_max_ = 2*π*NA*/nλ*_ex_ denotes the maximum lateral frequency support of the Optical Transfer Function (OTF) of the microscope’s objective with numerical aperture (NA) and operating at a wavelength of *λ*_ex_. For scenarios involving substantial spectral Stokes shifts, consult Ref. [14].

**Fig. 1.**
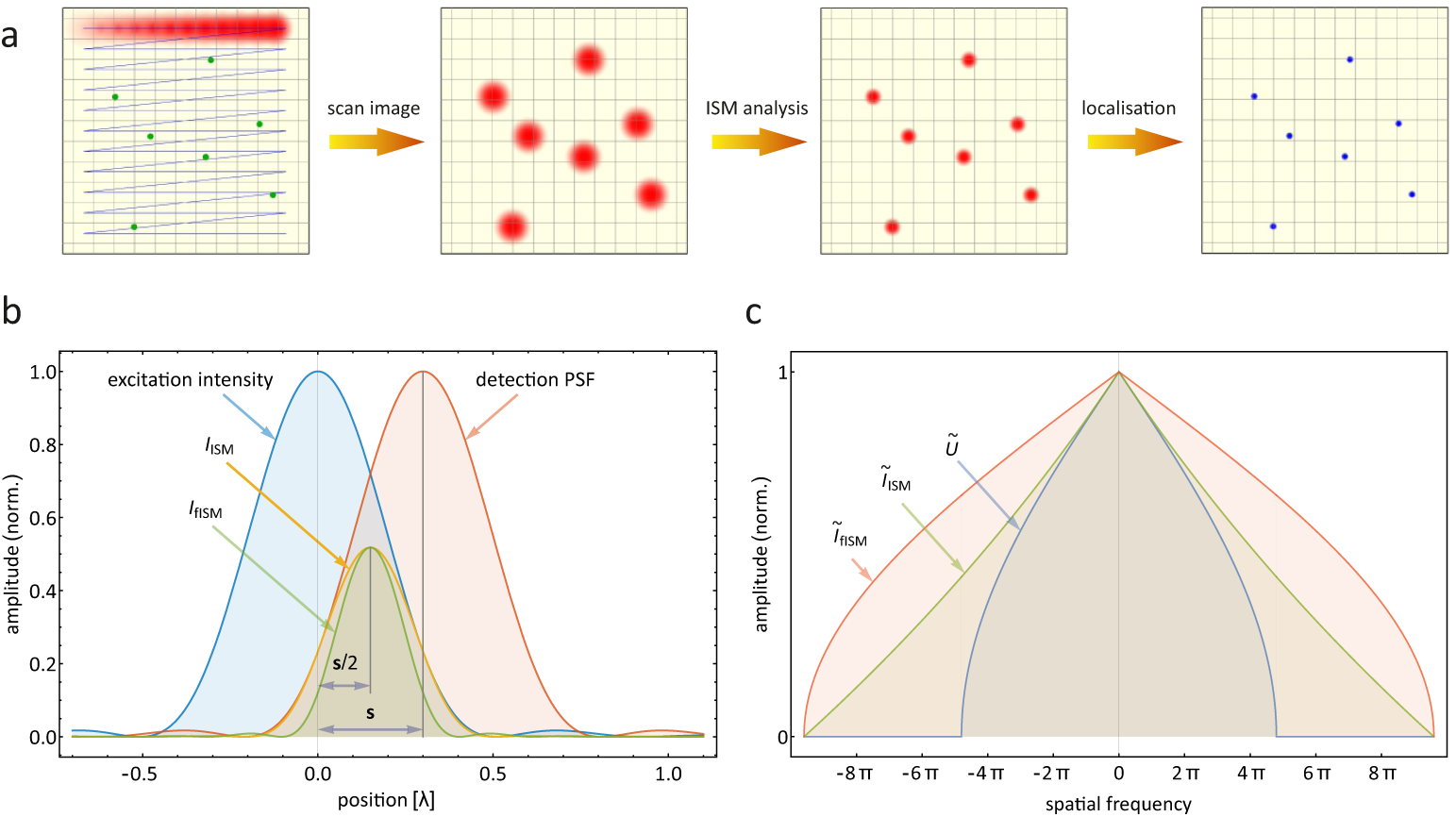
(a) Workflow of an iSMLM experiment, from left to right: images are recorded with a rapid confocal laser scanning microscope (left panel), which yield, for scarcely labeled samples, distributions of single molecule images (second from left panel). By applying an ISM algorithm to these images, the width of single molecule images is reduced by a maximum factor of 2 (third from left panel), which ideally yields a doubled localization accuracy (rightmost panel). (b) Principle of Pixel Reassignment an Image Scanning Microscopy: The curve ’excitation intensity’ illustrates a cross-section of the excitation intensity distribution, while the curve ’detection PSF’, centered at position **s**, represents the laterally shifted detection efficiency of a single detector pixel at position **s**. The resulting scan image of a single molecule, as captured by this particular pixel, exhibits a shape depicted by the curve *I*_ISM_, which is centered at **s***/*2. Subsequently, a single molecule image following Fourier reweighting is denoted as *I*_fISM_ and possesses precisely half the width of the excitation intensity and detection efficiency distributions. All calculations ere done for a water immersion objective with 1.2 NA and setting the refractive index of water to 1.33. The horizontal axis is in units of wavelength. (c) Optical Transfer Functions: The curve *Ũ* illustrates a cross-section of the Optical Transfer Function (OTF) associated with the excitation intensity distribution or wide-field imaging. The curve *Ĩ*_ISM_ represents the OTF for raw ISM data without Fourier reweighting. In contrast, the curve *Ĩ*_fISM_ portrays the ideal OTF achieved after Fourier reweighting. This reweighted OTF is essentially a re-scaled version of *Ũ*, with the horizontal axis expanded by a factor of two. The horizontal axis is in units of one over wavelength. The maximum value of all curves is normalized to one.

**Fig. 2.**
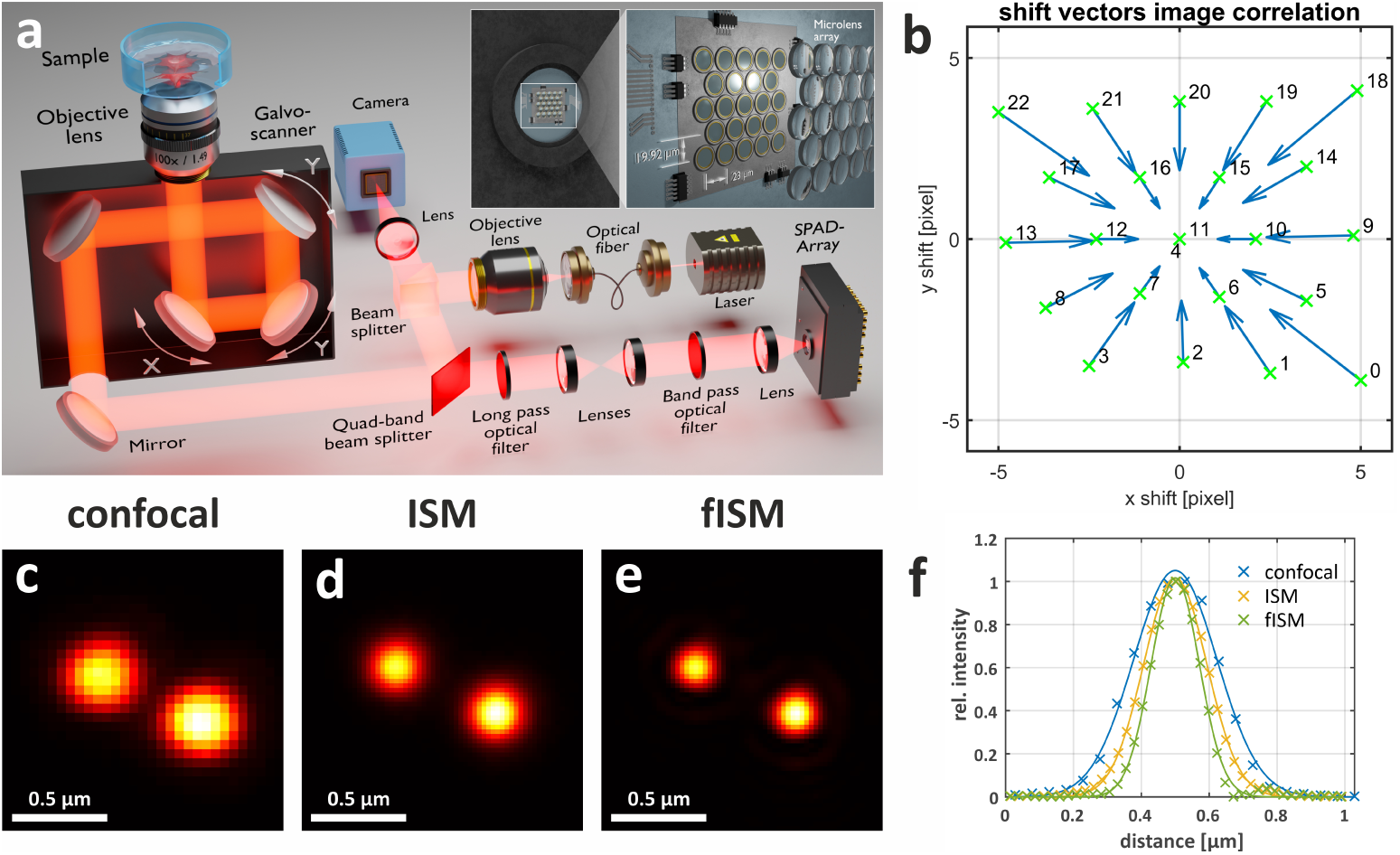
(a) Schematic of the FL-iSMLM setup. The inset depicts the arrangement of the 23 individual SPAD pixels and a micro lens array, which increases the fill factor of the detector up to 82%. (b) Shift vectors for ISM pixel reassignment. Green crosses represent pixel positions relative to the optical axis, and blue arrows indicate the ISM shift vectors. Pixel 4 does not have a shift vector because it is discarded due to its significantly increased dark-count rate (1.5 kcps at 7*^◦^*C), so its position in the panel overlaps with that of the central pixel 11. The average dark-count rate across the other 22 pixels is 35 cps at 7*^◦^*C. (c, d, and e) Images of sub-diffraction beads (Gatta Beads red, 23 nm, labeled with Atto 647N, GATTAQUANT GmbH). (c) Confocal image obtained by summing the signal over all detector pixels. (d) Photon detection positions adjusted according to ISM pixel reassignment. (e) Fourier-reweighted ISM image. The Pixel size in panel (d) and (e) has been doubled by Fourier transforming the image and padding the resulting image with zeros to double the size before transforming it back. (f) Intensity trace across one bead. The calculated 2*σ* resolution is 246*±*4 nm, 186*±*1 nm, and 142.2*±*1.7 nm for confocal, ISM, and Fourier-reweighted ISM, respectively.

A cross-section of a scan image of a single molecule recorded by a detector pixel at position **s** is depicted by the green curve centered at **s***/*2 in Fig. 1(a). It results from the product of the molecule’s excitation efficiency and the efficiency of detecting its fluorescence emission, as defined in Eq. 2. To obtain a final iSMLM (ISM) image, it is necessary to optimally combine the scan images captured by all detector pixels in such a way that they align with each other. Therefore, before adding a scan image recorded by a detector pixel at position **s** to the final ISM image, it should be shifted by **s***/*2 toward the center [12, 13]. Mathematically, this operation is described as follows:

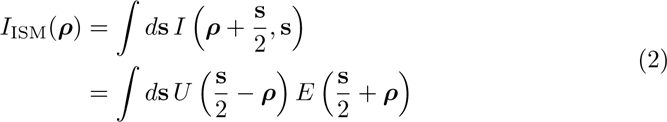

This shifting operation, also known as pixel reassignment [14, 31], involves shifting the intensity recorded by detector pixel **s** by **s***/*2 toward the center of the excitation focus for each scan position of the laser focus.

To comprehend why this reduces the size of a single-molecule image, consider the Fourier transform *Ĩ*_ISM_(**q**) of *I*_ISM_(***ρ***), where **q** is the Fourier variable corresponding to ***ρ***:

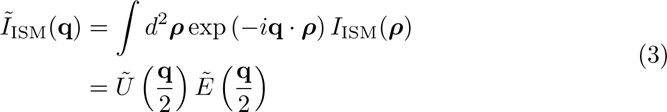

Cross-sections of the Optical Transfer Functions (OTFs) for excitation intensity distribution *Ẽ* (**q**) and wide-field detection *Ũ* (**q**) are shown by the blue curve in Fig. 1(b), denoted as *Ũ*. The product *Ẽ*(**q***/*2)*Ũ*(**q***/*2) is plotted as the green curve labeled *Ũ*_ISM_. It demonstrates that the lateral frequency support of *Ĩ*_ISM_(**q**) is twice as wide as that of the original OTFs *Ẽ* (**q***, z*) and *Ũ* (**q**). Thus, in principle, the image *Ĩ*_ISM_(***ρ***) exhibits double the resolution compared to an image captured by a pure laser-scanning or pure wide-field microscope. However, as demonstrated in Ref. [34], the optimal OTF of a wide-field microscope with a doubled frequency support is not given by Eq. 3 but by the original OTF rescaled by a factor 1/2, i.e. by *Ũ* (**q***/*2). This rescaled OTF is shown by the red curve in Fig. 1(b) and labeled with *Ĩ*_fISM_. Thus, to achieve the full resolution enhancement offered by ISM, the spatial frequencies in *Ĩ*_ISM_(**q**) should be reweighted with the weight function *Ũ* (**q***/*2) */Ĩ*_ISM_(**q**) to approach *Ũ* (**q***/*2).

However, as seen in Fig. 1(b), the function *Ĩ*_ISM_ approaches zero when the length of the wave vector approaches the limits of the frequency support, so that division by *Ĩ*_ISM_ which will lead to amplification of noise for large frequencies. To avoid this, one can introduce a regularizing factor *ϵ* so that the weight function reads

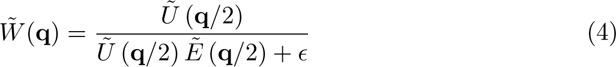

instead of *Ũ* (**q***/*2) */Ĩ*_ISM_(**q**). Usually, *ϵ* can be chosen as small as *∼* 0.1 of the maximum amplitude of *Ũ* (**q***/*2) *Ẽ* (**q***/*2), so that potential nonlinearities introduced by this procedure are kept small. Certainly, there exist more advanced regularization methods such as Wiener filtering (which takes image noise properties into account, see e.g., [35]), but for the sake of simplicity, we have focused here on the simple Fourier reweighting with weight function of Eq. 4. Thus, the Fourier transform of a single-molecule image is approximately:

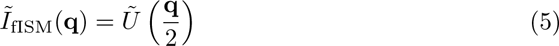

This Fourier transform corresponds to that of an image of a molecule captured with a wide-field microscope with doubled resolution.

In the final step of iSMLM, the Fourier-reweighted ISM images of single molecules are processed using standard single-molecule localization software [36–39] to obtain a final SMLM image. Starting with single-molecule images half as wide as those in conventional wide-field SMLM, the final localization accuracy and image resolution are doubled. Alternatively, iSMLM achieves roughly the same resolution as conventional SMLM with only a quarter of the photon budget [32].

### 2.2 Super-resolution imaging with iSMLM

#### 2.2.1 FL-iSMLM Measurements

To showcase the localization improvement achieved by iSMLM, we immunolabeled *α*-tubulin in COS-7 cells using Alexa 647-conjugated antibodies. The on-off photoswitching of the fluorophores was initiated using a PBS buffer containing cysteamine (see also section ’Imaging buffers’ in Supplementary Information). Fine-tuning the blinking behavior, and consequently the number of photons per localization event, was achieved by varying the composition of this photoswitching buffer. For the dSTORM measurement, we initially selected a suitable region of interest where most of the microtubules lay in the same plane. Subsequently, we defined a 5 µm *×* 5 µm region of interest, as shown in zoom-in I. Images were captured at a frame rate of 27.9 Hz. Approximately 5 *·* 10^4^ to 7.5 *·* 10^4^ images were recorded to ensure a sufficient number of single molecule blinking events.

To optimize single-molecule localization, we applied frame binning by combining 10 frames before conducting further image analysis. This approach helps prevent PSF distortions arising from photoswitching during scanning, ensuring an even distribution of the photon budget across all scan positions.

For localization and lifetime analysis, we employed the open-source tool TrackNTrace [38], which was extended to include ISM pixel reassignment, adjusting the position of each recorded photon event during the measurement [40]. Subsequently, Fourier reweighting was applied to the resulting ISM images, following the method described in Ref. [13] (as detailed in section ’Data analysis’ below).

The resulting images were then used for single molecule localization, ultimately yielding the final super-resolved image. Fig. 3(b)-(d) shows the images reconstructed using three different approaches: dSTORM without ISM photon reassignment, with ISM photon reassignment, and with photon reassignment plus Fourier reweighting, respectively. To determine the lifetime of each localized single molecule, we collected all photons from the event and performed tail-fitting using a mono-exponential decay function on the corresponding TCSPC histogram (see section ’Data analysis’ above). Notably, photons from molecules appearing in subsequent frames were combined into a single event.

**Fig. 3.**
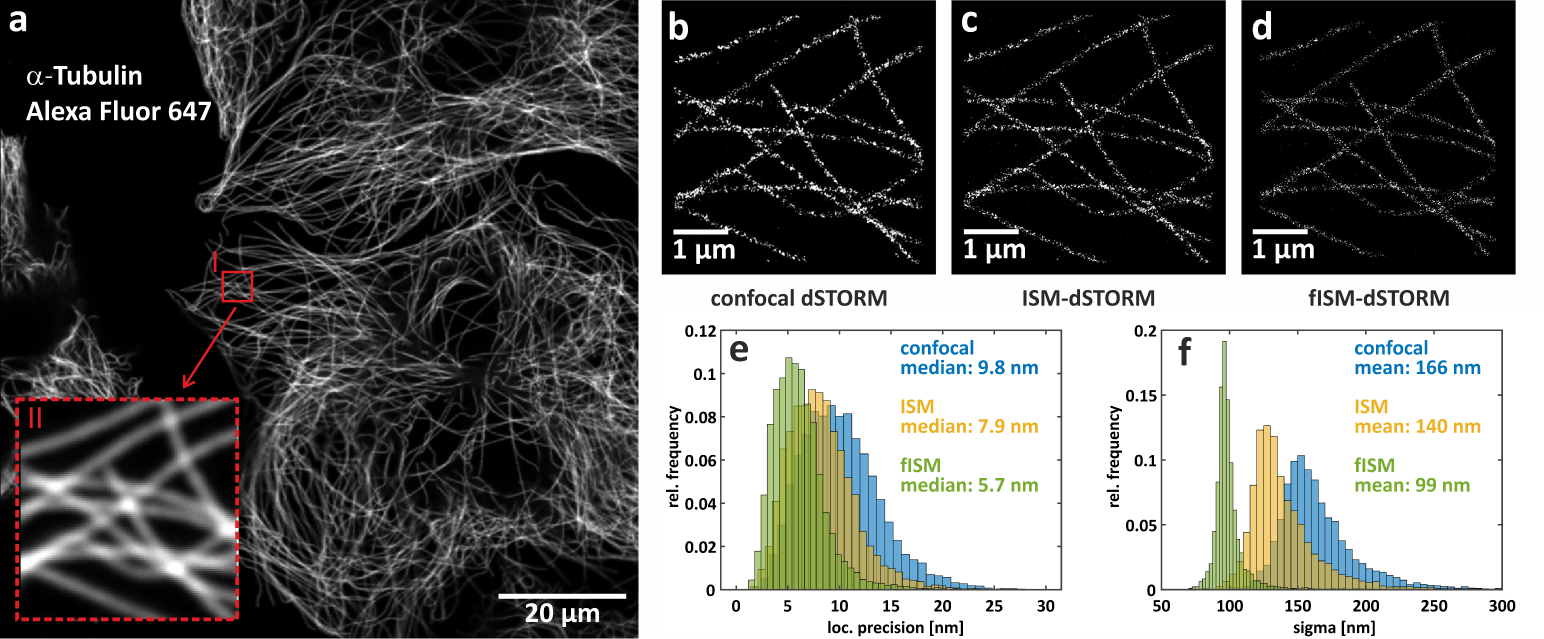
Single labeling iSMLM dSTORM results. (a) Overview of a COS-7 cell where *α*-tubulin was labeled with Alexa 647. Zoom I shows a confocal image of the actual 5 µm *×* 5 µm region of interest used for dSTORM analysis.(b-d) Confocal, ISM and Fourier-reweighted ISM dSTORM reconstruction from 5 *·* 10^4^ images with 10 frame binning. (e) Localization precision for confocal, ISM and Fourier-reweighted ISM dSTORM of *α*-tubulin. (f) Histograms of fitted *σ* values of localization events for confocal, ISM and Fourier-reweighted ISM dSTORM.

To enhance the fidelity of the image reconstruction and eliminate localizations that could result from single molecule mis-identifications, we only considered events with more than 200 photons. This number has been chosen for all three reconstruction methods. The selection was made after analyzing the relative enhancement of localization precision between confocal SLMSM and fISM SMLM as a function of minimum number of photons. This had shown a stark dependency at low number of photons, but a plateau for filtering 200 or more, as can be seen in SI Fig. 2. The width of a single molecule image was defined as the 2*σ* size of a fitted circularly symmetric 2D Gaussian intensity distribution. Additionally, molecule images that occurred in several subsequent frames were combined into a single event.

The average *σ* value of single molecule images (the average *σ* values of Gaussian fits) for the single-molecule localizations in Fig. 3(b) was 166 nm, as indicated in panel (f), which falls within the range of the measured PSF determined from the bead calibration measurements. The median localization precision of dSTORM was estimated at 9.7 nm, as calculated using a modified Mortensen’s equation [41]. This value improved to 7.9 nm after ISM photon reassignment, with a corresponding *σ* value of 140 nm. The reconstructed image, Fig. 3(c), also revealed finer details, particularly noticeable in regions where microtubules overlap. Finally, after Fourier reweighting, the localization precision further increased to 5.7 nm. The improvement is also shown by a more pronounced separation of the two peaks in the microtubule cross-section, depicted in Fig. S1. We observed that the mean peak-to-peak distance was 41 *±* 2 nm, including the linkage error, which is consistent with the literature results presented earlier [42].

#### 2.2.2 FL-iSMLM Multiplexing

The availability of lifetime information in FL-iSMLM allows for image multiplexing, enabling the simultaneous imaging of different targets based on the distinct fluorescence lifetimes of the applied labels. To demonstrate this capability, we labeled COS-7 cells with *α*-tubulin and F-actin using antibody-conjugated Alexa 647 and phalloidin-Janelia Fluor 646, respectively (for sample preparation and imaging buffers, see Supporting Information). Both fluorophores exhibited comparable on-off photoswitching behavior and similar quantum yields under the same buffer conditions, making them an ideal combination for lifetime-multiplexed FL-iSMLM.

In Fig. 4(a), we present a confocal image of one of these cells, where color represents the average fluorescence lifetime, calculated as the mean photon arrival time. The square denotes the region of interest chosen for dSTORM measurement. In this analysis, we recorded 6.5 *·* 10^4^ images at a frame rate of 27.2 Hz, totaling just under 40 minutes of acquisition time. The resulting data underwent analysis using our open-source software, TrackNTrace, with the newly incorporated ISM plugin [40].

**Fig. 4.**
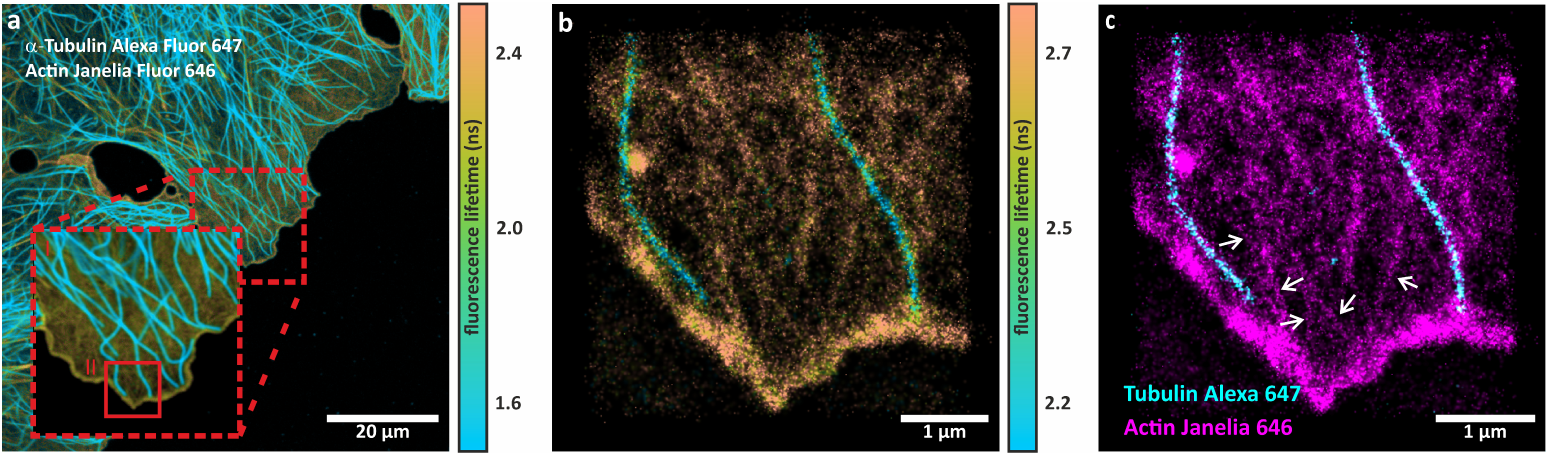
dSTORM based FL-iSMLM multiplexing. (a) Overview of a COS-7 cell where *α*-tubulin was labeled with Alexa 647 and actin with phalloidin-Janelia Fluor 646. Zoom II shows the actual 5 µm *×* 5 µm region used for dSTORM analysis. (b) Super-resolved FL-iSMLM dSTORM reconstruction from 6.5 *·* 10^4^ images with 10 frame binning. (c) False color super-resolved image based on fluorescence lifetime data. The two lifetime components were separated using our maximum-likelihood based lifetime-pattern matching classifier. Localization events identified as Alexa 647 are colored cyan, and localization events identified as Janelia Fluor 646 are colored magenta. White arrows indicated a visible actin filaments.

For each localization event, we determined the fluorescence lifetime by collating all photons associated with the event and subsequently performing a mono-exponential tail-fitting using a Maximum Likelihood Estimator (MLE). Tail-fitting involves removing the excitation peak from the TCSPC curve and retaining only photons arriving later than 0.3 ns after the TCSPC curve’s peak. The resulting FL-iSMLM superresolution image is displayed in Fig. 4(b), revealing a distinct separation of structures based on lifetime: The short lifetime events at *∼*1.7 ns correspond to Alexa 647-labeled *α*-tubulin, while the long lifetime components at *∼*2.8 ns correspond to actin filaments. We also employed a maximum-likelihood based pattern-matching classification [43] to distinguish these two structures. This involved first determining the decay patterns for only Alexa 647-labeled *α*-tubulin and phalloidin-Janelia Fluor 646-labeled actin using samples with only one respective labeling. We then utilized these patterns to separate the single-molecule events in the doubly-labeled samples. The resulting false color image is presented in Fig. 4(c), with Alexa 647 localizations represented in cyan and Janelia Fluor 646 localizations in magenta. In this context, localizations with a probability of less than 99% of corresponding to their respective decay patterns were systematically excluded from the reconstruction. This probability was determined by calculating the posterior probability of each single molecule’s TCSPC curve belonging to one of the reference decays [43]. Notably, the FL-iSMLM image unveils intricate actin filament structures that were previously indiscernible in the confocal image. Although actin filaments are not as evident as microtubules due to several factors related to the actin network properties e.g. label density (microtubules have about six times more binding sites per unit length than actin) [44], we were able to resolve thin actin filaments (see white arrows in Fig. 4(c)) even in a simultaneous multi-target experiment and with a non-standard fluorescence dye.

We conducted a repeat dSTORM measurement at the center of another COS-7 cell, which exhibited a comparable density of actin and tubulin. An overview of this cell is presented in Fig. S3(a), where color is used to represent fluorescence lifetime. In Fig. S3(b), we showcase a super-resolved fluorescence-lifetime image obtained through FL-iSMLM within the region enclosed by the red square. The corresponding lifetime histogram reveals two distinct and well-separated peaks, both of which have been fitted with Gaussian distributions.

To generate the dual-color image featured in Fig. S3(d), we once again employed a maximum-likelihood based pattern-matching classifier. This classifier utilized the same pre-established decay patterns as those applied in Fig. 4.

To showcase the capabilities of FL-iSMLM on a biologically more relevant sample, we employed DNA-PAINT [45] to visualize pre- and postsynaptic scaffolds, Bassoon and Homer1, within cultured hippocampal neurons. These proteins are situated across the synaptic cleft and are known to form domains that organize into relatively flat structures: the presynaptic active zone and the postsynaptic density (PSD), respectively. To label these proteins, we used specific antibodies conjugated with DNA target strands. Bassoon was imaged using DNA imager strands labeled with Cy3b, while Homer1 was imaged using DNA imager strands labeled with Atto 565. For details on cell cultures, sample preparation, and imaging buffers, see Supporting Information.

In Fig. 5(a), we present FL-iSMLM reconstructions of a 7 µm *×* 7 µm field of view in a hippocampal cell. Confocal scan images were acquired with 70 nm scan steps and the same dwell time as in previous measurements. We recorded a total of 10^5^ images over an acquisition time of just over one hour. To account for the DNA-PAINT binding rates, we increased frame binning from 10 to 20 frames. Image reconstruction was once again performed using TrackNTrace, which was amended with ISM capabilities [40]. The localization events were classified using fluorescence lifetime maximum-likelihood based pattern matching, following the same methodology as previously described.

**Fig. 5.**
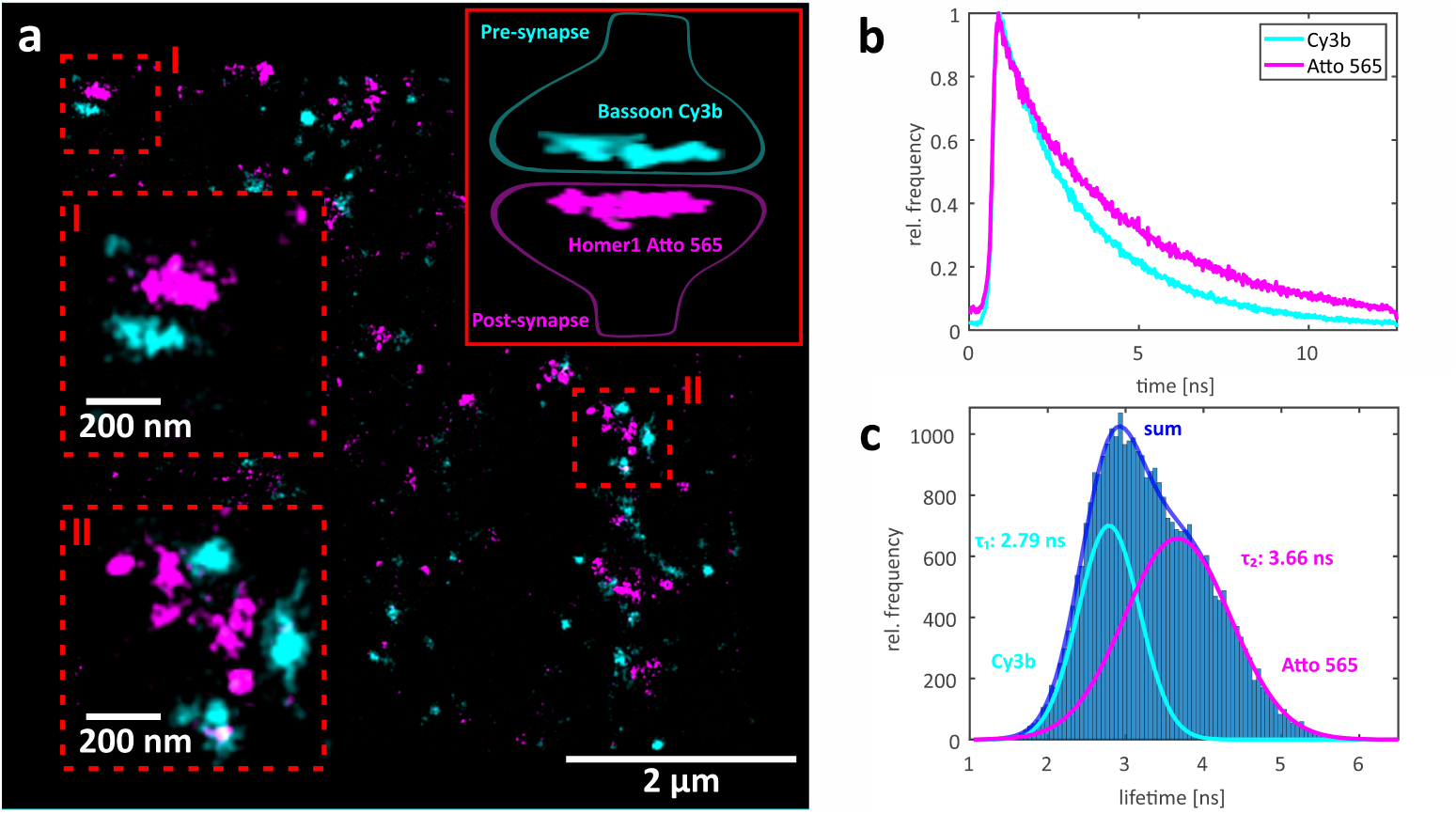
DNA-PAINT-based FL-iSMLM multiplexing. (a) Super-resolved Fourier-reweighted iSMLM image of Homer1 labeled with Atto 565 (magenta) and Bassoon labeled with Cy3b (cyan). Targets were separated by using a maximum-likelihood based fluorescence-lifetime pattern-matching classification. The schematic illustration of the pre-synapse and post-synapse is presented in the upper corner. (b) Fluorescence lifetime decay curves for pure Cy3b and Atto 565 samples, respectively. (c) Fluorescence lifetime histogram obtained from single molecule localizations in (a), fitted with two Gaussian distributions.

For this analysis, reference lifetime patterns were derived from DNA-PAINT fluorescence lifetime images of pure samples in which either Bassoon or Homer1 was exclusively labeled. The resulting TCSPC histograms are displayed in Fig. 5(b). Notably, we observed a subtle lifetime difference of only 0.9 ns between Cy3B and Atto 565 labeled proteins, as depicted in Fig. 5(c). This minor difference necessitated the use of pattern matching for accurate lifetime classification of single molecule localization events. In the reconstructed images, localization events attributed to the decay pattern of Cy3B, with a probability exceeding 99%, are depicted in cyan, whereas those associated with Atto 565, with an equivalent confidence level, are indicated in magenta. All other events have been excluded from this reconstruction.

## 3 Discussion

In this study, we introduced a novel concept in SMLM by integrating it with ISM. This integration leads to a remarkable enhancement in resolution, up to a factor of two, or equivalently, a reduction in the required photon budget for a given resolution by up to four times. A recent theoretical study [46] even suggests that, with advanced data analysis techniques, the resolution enhancement achievable with iSMLM could surpass the factor of two.

To implement ISM, we incorporated a SPAD array detector into a CLSM. Coupled with a high-speed TCSPC electronics, this upgraded CLSM not only enhanced the precision of molecular localization but also enabled the capture of fluorescence lifetimes of individual molecules. Moreover, CLSM provides the advantage of confocal sectioning and limited photobleaching due to the fact that only the region of interest is exposed.

We ensured the improvement in lateral resolution had been achieved through our implementation of ISM by imaging sub-diffraction fluorescent beads. The attained increase by a factor of 1.73 is comparable to other publications implementing Image Scanning Microscopy [13, 27]. The main reasons for the discrepancy between the theoretically possible increase by a factor of two are the Stokes shift between excitation and emission light, a non-zero background caused by the dark count of the detector, as well as stray light. This background compelled us to increase the regularization parameter during Fourier-reweighting to prevent noise amplification, limiting the maximum spatial frequency. Our initial measurements of microtubules, presented in Fig. 3 excellently demonstrated that this resolution enhancement could also increase the localization precision of SMLM from 9.8 nm to 5.7 nm, corresponding to an improvement by a factor of 1.72. The slight deviation from the expected 1.73 value could have multiple causes. Firstly, the measurement conditions are vastly different during the iSMLM recordings because of the low signal-to-noise levels, compared to the fluorescence bead measurements. Nonetheless, in order to enhance the coherence of our assertions, no additional measures were implemented during the analysis of the data. Such as median filtering or noise reduction. Our simulations have shown that background noise can play a major role in iSMLM. Secondly, the axial position of the microtubules does vary in contrast to the fluorescence bead, which all lay in the same plane. This capability opened the door to multi-target SMLM without the limitations of chromatic aberration, which is crucial when aiming to co-localize structures on the nanometer length scale.

To promote wider adoption of iSMLM, we extended the capabilities of our publicly available single-molecule tracking and localization software, TrackNTrace [38]. We also introduced a novel FL-ISM-plugin [40], harnessing the unique lifetime multiplexing feature of our FL-iSMLM.

With FL-iSMLM’s distinctive features, we simultaneously visualized two essential components of the cytoskeleton: microtubules and actin filaments. Even in scenarios with significant differences in target protein concentrations, our highly efficient maximum-likelihood based pattern matching classification reliably distinguished and separated both structures, without the need to eliminate a significant number of localizations. This is easily verified by comparing Fig. 4 (b) and (c), where one can see that almost all single molecule events are present in both reconstructions. In particular, Fig. 4 (b) was reconstructed from 48225 and Fig. 4 (c) from 47621 localizations, which corresponds to discarding only 1.24% of all events. In comparison, the percentage of 2.25% of events discarded by the applied photon number threshold is about twice as large. Furthermore, we employed DNA-PAINT-based FL-iSMLM to achieve nanometer-scale precision in depicting the spatial arrangement of synaptic scaffolds, specifically Homer1 and Bassoon. These proteins form closely apposed structures on opposite sides of the synapse, and our technique allowed us to visualize their arrangement with exceptional precision. The average distance between Bassoon and Homer1 across the synaptic cleft, determined by measuring the separation between the peak positions of their respective localizations (as exemplified in Fig. S4), was measured at *∼* 150 nm. This measurement is consistent with previously published values [47, 48]. It should be mentioned that due to the much smaller lifetime difference between the Cy3b and Atto 565 dyes, 8502 of all 40945 localizations were discarded, corresponding to 20.8%.

The major limitations of this novel technique are the frame rate and field of view size. Both are inherent to all scanning-based microscopy systems and are unavoidable when combining SMLM with Image Scanning Microscopy. On one hand, a certain pixel size is required to adequately sample the PSF of each single molecule. In general, this size cannot exceed more than 80 nm for ISM. On the other hand, a frame rate between 20 Hz and 30 Hz is required to capture single events accurately in dSTORM or DNA-PAINT, as explained before. This requirement sets an upper limit to the image size at around 100 by 100 pixels due to the limited dwell time of our confocal scanner. Consequently, this results in a field of view of not more than 8 µm by 8 µm, taking into account the corresponding frame rates and resulting in approximately 30 minutes of measurement time.

In conclusion, iSMLM represents a significant expansion of ISM into the realm of SMLM, akin to ISM extensions of other super-resolution modalities such as STED microscopy [49], or MINFLUX [50]. When compared to STED or MINFLUX, iSMLM distinguishes itself with its simplicity, versatility, and seamless incorporation of fluorescence lifetime imaging capabilities. We posit that FL-iSMLM, offering doubled resolution and eliminating chromatic aberration when using fluorescence lifetime for multiplexing, establishes a groundbreaking benchmark in SMLM. This achievement gains paramount significance as SMLM approaches the molecular length scale in its pursuit of higher resolution.

## Methods

### Setup

ISMLM in this study was realized with a custom-made ISM setup illustrated in Fig. 2(a). For excitation, we employed a 640 nm pulsed laser diode with a 40 MHz repetition rate and a pulse width of *≤*90 ps (PDL 800-B driver with LDH-D-C-640 diode, PicoQuant) and a 561 nm supercontinuum white light laser with a 80 MHz repetition rate (Fianium WhiteLaser SC450, NKT Photonics). The laser beam was coupled into a single-mode optical fiber (PMC-460Si-3.0-NA012-3APC-150-P, Schäfter + Kirchhoff) using a fiber coupler (60SMS-1-4-RGBV-11-47, Schäfter + Kirchhoff).

After the fiber, the laser light was collimated using an air objective (UPlanSApo 10x/0.40 NA, Olympus) and passed through a clean-up filter (MaxDiode 640/8, Semrock or BrightLine 563/9, Semrock) before being directed to the microscope through an ultra-flat quad-band dichroic mirror (ZT 405/488/561/640 RPC, Semrock). Laser scanning was achieved using a three-galvo mirror scan system (FLIMbee, PicoQuant) with an integrated 2x magnification telescope. The laser light was focused onto the sample using a high numerical-aperture oil immersion objective (UApo N 100×/1.49 NA oil, Olympus).

Samples were mounted on a manual xy-stage (Olympus) equipped with a z-piezo stage (Nano ZL100, Mad City Labs) for precise focus control. The same objective was used for collecting the fluorescence light (epi-fluorescence configuration), which was then de-scanned in the FLIMbee system. After passing through the quad-band beam splitter, the emission light was focused through a pinhole (100-µm P100S, Thorlabs) using an achromatic lens (TTL180-A, Thorlabs). The pinhole was solely used for optical component alignment and was removed during the ISM and iSMLM measurements.

To eliminate back-reflected and scattered light, a long-pass filter (635 LP or 561 LP Edge Basic, Semrock) was placed in front of two achromatic lenses (AC245-100-AML and AC254-075-A-ML, Thorlabs), which relayed the emission light onto a SPAD array detector (preliminary version of PDA23, PicoQuant). The SPAD array consists of 23 individual SPADs arranged in a hexagonal pattern, with a pixel-to-pixel distance of 23 µm. This array is equipped with a matching microlens array, which boosts its fill factor beyond 80%, leading to a total photon detection efficiency of 23.5% at 700 nm and 35% at 600 nm. A band-pass filter (BrightLine HC 708/75, Semrock or BrightLine HC 617/73, Semrock) was employed to further suppress scattered light.

The output signals from the 23 pixels were independently recorded using a TCSPC module (Multi Harp 160, PicoQuant), controlled by the SymPhoTime 64 software (PicoQuant). The software also controlled the galvo and piezo scanners. Samples were scanned with step sizes of 50 nm to 70 nm, with a dwell time of 2.5 µs and a temporal resolution for photon-detection event timing of 20 ps.

### Calibration

ISM calibration and SMLM measurements were conducted using our custom-made time-resolved laser scanning microscope, as depicted in Fig. 2(a) for detailed specifications. This microscope is equipped with a high-speed galvo scanner capable of capturing scan images at high speeds.

For excitation, we employed either a 640 nm or 561 nm pulsed laser. This setup, combined with our picosecond multichannel event timer, enabled us to utilize TCSPC for precise fluorescence lifetime measurements.

The analysis of the acquired data was carried out using our in-house software, TrackNTrace [40], specifically amended for this purpose.

To compute the correct shift vectors for ISM pixel reassignment (similar to the phase correlation approach used in [27]) and validate the resolution improvement, we conducted scans of sub-diffraction beads under the same conditions as used for the ISM dSTORM and DNA-PAINT measurements. Fig. 2(b) illustrates the calculated shift vectors.

To determine these vectors, we employed image correlation to ascertain the shift of each individual detector pixel relative to the optical axis. Prior to this, the SPAD array was aligned so that the central pixel of the hexagonal pattern (as seen in Fig. 2(a)) coincided with the optical axis. Subsequently, the scan images captured by the various individual pixels of the SPAD array were correlated with the scan image recorded by the central pixel. The position of the maximum in the resulting correlation map provided the necessary shift for the scan image recorded by a specific pixel in relation to the central pixel. These reassignment vectors are illustrated in Fig. 2(b), where the green crosses represent the actual positions of the 23 pixels, and the arrows indicate the direction and magnitude of the shift. The units are given in the number of scan steps (in our case, 50 nm).

Fig. 2(c)-(e) demonstrate the resolution enhancement of ISM and Fourier-reweighted ISM compared to Confocal Laser Scanning Microscopy (CLSM). To quantify this improvement, we plotted intensity profiles across one of the beads and calculated its 2*σ* width, as shown in Fig. 2(f). The confocal image, obtained by summing photons from all detector pixels, yielded a 2*σ* value of 248 *±* 4 nm. With ISM reassignment, this value decreased to 186 *±* 1 nm. Further Fourier reweighting, as detailed in [13], improved this value to 142.2*±*1.7 nm, resulting in a total improvement by a factor of 1.73 compared to CLSM.

### Data analysis

FL-iSMLM measurements were analyzed using an extended version of the TrackNTrace software package [40]. To generate images from the raw scan data, we combined 10 or 20 scans into a single frame(frame binning). In the localization process, we utilized the ”Cross-Correlation” detection plugin with default parameters and the ”TNT Fitter” refinement plugin, employing pixel-integrated Gaussian Maximum Likelihood Estimation (MLE) fitting.

Localizations found in adjacent frames with a separation of less than 100 nm were considered to originate from the same molecule. Their positions were subsequently re-fitted using the photon detection events from all relevant frames. For the purpose of lifetime fitting, we constructed a TCSPC histogram for each localized molecule by collecting all photons within a circle with a radius of less than 2 times the standard deviation (*σ*_PSF_) from the molecule’s center position. These TCSPC histograms were then fitted with a mono-exponential decay function using Maximum Likelihood Estimation (MLE) to determine the lifetime, as detailed in Ref. [43].

In the context of iSMLM, photon positions were reassigned utilizing ISM shift vectors obtained through the imaging of PSF beads with a diameter of 23 nm, as discussed in the preceding section. This reassignment was performed prior to frame binning and event localization. Fourier reweighting was applied to the combined frame data following the algorithm detailed in Ref. [13]. The necessary OTF was computed through the Fourier transformation of the detection PSF from our microscope, which had been parameterized using the same bead image employed for shift vector calibration. For the normalization parameter *ɛ* (see Eq. 4 and Ref. [13]), which averts the amplification of image noise at high frequencies in Fourier space, we selected a value of 0.1 of the maximum of the OTF. This number has been chosen to be the smallest value where ring formation around a single emitter due to noise amplification does not occur. For *ɛ* = 0 this method would have provided the theoretically optimal way to increase image resolution for our system [34]. Both photon reassignment and Fourier reweighting have been incorporated into the extended version of TrackNTrace [40].

### Cell culture and immunostaining

#### COS-7 cell line culture

A COS-7 African Green Monkey fibroblast-like cell line was cultured at 37*^◦^*C with 5% CO_2_ in T25-culture flasks, using Dulbecco’s Modified Eagle Medium (DMEM) supplemented with 10% FBS (Sigma-Aldrich, F7524) and 100 U/ml penicillin + 0.1 mg/ml streptomycin (Sigma P4333). One day before use, cells were seeded onto 35 mm glass bottom Ibidi dishes (µ-Dish 35 mm high, 81158) at a concentration of 10^5^ cells/dish, and cultivated overnight.

#### Hippocampal cultured neuron

Experimental procedures involving animals (Wistar rats, P0 to P1) were conducted in accordance with the guidelines of the local regulatory authority, the Lower Saxony State Office for Consumer Protection and Food Safety (Niedersächsisches Landesamt für Verbraucherschutz und Lebensmittelsicherheit), under license Tötungsversuch T09/08. Briefly, hippocampi were dissected from rat brains, followed by washing using Hank’s Balanced Salt Solution (HBSS, 14175053, Invitrogen, Waltham, MA, USA). Then, hippocampi were gently agitated in a digestion solution, which included 15 U/ml papain (LS003126, Worthington, Lakewood, USA), alongside with 1 mM CaCl_2_, 0.5 mM EDTA, and 0.5 mg/ml L-cysteine (30090, Merck), all dissolved in DMEM. This enzymatic digestion was performed for 1 h at 37*^◦^*C, before inactivation with a buffer solution containing 10% FCS and 5 mg/ml bovine serum albumin (BSA, A1391, Applichem, Darmstadt, Germany) in DMEM. After 15 minutes, this inactivation solution was replaced with growth medium containing 10% horse serum (S900-500, VWR International GmbH, Darmstadt, Germany), 1.8 mM glutamine, and 0.6 mg/ml glucose in MEM (51200046, ThermoFisher Scientific), which was utilized for the subsequent iterative washes of the hippocampal tissue. Afterwards, the neurons were isolated by trituration, a method that uses a glass pipette for mechanical disruption, then separated by centrifugation at 154 ×g (8 minutes), and re-suspended in the same medium. The neurons were then seeded onto poly-L-lysine-coated coverslips and allowed to adhere for several hours before being transferred to Neurobasal™-A medium (10888022, ThermoFisher Scientific) with 0.2% B27 supplement (17504-044, ThermoFisher Scientific) and 2 mM GlutaMAX (35050-038, ThermoFisher Scientific). Finally, neurons were maintained in a humidified incubator (5% CO_2_, 37*^◦^*C) for at least 14 days before use.

#### Immunostaining of cytoskeletal proteins

For microtubule and actin immunolabeling, COS-7 cells were washed with pre-warmed (37*^◦^*C) PBS, and permeabilized for 2 min with 0.3% glutaraldehyde (GA) + 0.25% Triton X-100 (Sigma-Aldrich, 340855 and Thermo Fisher, 28314) in pre-warmed (37*^◦^*C) cytoskeleton buffer (CB) at pH 6.1, consisting of 10 mM MES (Sigma-Aldrich, M8250), 150 mM NaCl (Sigma-Aldrich, 55886), 5 mM EGTA (Sigma-Aldrich, 03777), 5 mM glucose (Sigma-Aldrich, G7021), and 5 mM MgCl_2_ (Sigma-Aldrich, M9272). After permeabilization, cells were fixed with a pre-warmed (37*^◦^*C) solution of 2% GA in CB for 10 min. After fixation, cells were washed twice with PBS and reduced with 0.1% sodium borohydride (Sigma-Aldrich, 71320) in PBS for 7 min. Cells were again washed three times with PBS before blocking with 5% BSA (Roth, 3737.3) in PBS for 1 h. Subsequently, cells were incubated with 10 ng/µl anti-alpha tubulin antibody (DM1A) (Abcam, ab264493) in a blocking buffer for 1 h. After primary antibody incubation, cells were washed thrice with 0.1% Tween 20 (Thermo Fisher, 28320) in PBS for 15 min. After washing, cells were incubated in a blocking buffer with 20 ng/µl of commercial secondary antibody goat anti-mouse IgG H&L (Alexa Fluor 647) (abcam, ab150115) for 45 min. After secondary antibody incubation, cells were again washed three times with 0.1% Tween 20 in PBS for 15 min. After washing, a post-fix with 4% formaldehyde (Sigma-Aldrich, F8775) in PBS for 10 min was performed, followed by three additional washing steps with PBS.

If actin was also fluorescently labeled, a final incubation step with a solution of 500 nM phalloidin-Janelia Fluor 646 in PBS for one hour was performed prior to imaging. This was followed by three washing cycles with PBS.

#### Immunostaining of synaptic proteins

Neurons were fixed with 4% PFA and 0.2% GA in PBS for 30 min before being quenched with a 0.1% sodium borohydride solution for 9 min. These cells were then exposed to a blocking/permeabilizing solution containing 5% BSA and 0.1% Triton X-100 in PBS for 30 min. Primary antibodies against Homer1 (160003, Synaptic Systems) and Bassoon (ADI-VAM-PS003-F, Enzo, New York, USA) were diluted in 5% BSA in PBS and applied to the coverslips for 60 min. Then, samples were washed three times with a solution of 0.1% Tween in PBS, followed by immersing the coverslips for 45 minutes in a diluted solution of secondary single domain antibodies with DNA strands (FluoTag^®^-XM-QC Anti-Mouse IgG kappa light chain + FAST docking site F1 and FluoTag^®^-XM-QC Anti-Rabbit IgG + FAST docking site F2) in 5% BSA in PBS. Samples were then washed three times with 0.1% Tween in PBS before imaging.

### Imaging buffers

For confocal dSTORM imaging with only Alexa 647 dye, a switching buffer consisting of 50 mM cysteamine in PBS at pH 7.4 was used. For dual-color imaging with both Alexa 647 and Janelia Fluor 646 dyes, an enzymatic oxygen scavenging buffer (glucose oxidase 0.5 mg/ml, catalase 40 g/ml, glucose 10% w/v in PBS at pH 7.4) with the addition of 50 mM cysteamine was used. For DNA-PAINT imaging of proteins in hippocampal neurons, the commercial imaging buffer MASSIVE-SDAB-FAST 2-PLEX kit (Massive Photonics) containing 770 pM Cy3b-labeled imager F1 and 500 pM Atto 565-labeled imager F2 was used.

### iSMLM simulations

We simulated single molecules with different photon and background levels to evaluate our findings and show how the different conditions in SMLM can have an effect on our measurements. For each combination of background and number of photons, we generated the PSF of a single point emitter using the scalar approximation [33] for each pixel of our SPAD-Array detector. To get the corresponding confocal PSF, the intensities of each detector pixel were summed up. For ISM the individual PSFs from the detector pixels were shifted according to the ISM reassignment vectors and summed up. In the final step, we used the same Fourier reweighting as described in the Data analysis section, with the same normalization parameter *ɛ* = 0.1 to create the fISM PSF. From this, we generated 1000 images of a single emitter for a given number of photos and signal-to-noise ratio. To simulate realistic photon emission, the three PSFs were normalized and the intensity in each pixel was drawn from a Poisson distribution with a rate *λ* equal to the product of the PSF value at the corresponding pixel and the number of photos. Background noise was added by drawing from a Poisson distribution with the same rate for all pixels. This rate was selected to fit the desired SNR. The resulting localization precision calculated by or single particle tracking software TrackNTrace and is plotted in images (a) to (c) of Fig. S2. Where each pixel corresponds to a combination of number of photons and signal-to-noise ration, and the colormap depicts the localization precision. Panel (d) shows the simulated improvement in localization precision between confocal SMLM and fISM SMLM in or iSMLM in blue and the measured in red. It is noteworthy that the normalization parameter selected for the simulation is identical to that used during the measurements, resulting in an improvement of less than two.

## Data Availability

All raw data is available in the following repository: https://gitlab.gwdg.de/igregor/s ingle-molecule-localization-microscopy-with-image-scanning-microscopy.git

## Code Availability

Every scrip and code to process the raw data is available in the following repository: https://gitlab.gwdg.de/igregor/single-molecule-localization-microscopy-with-image-scanning-microscopy.git

## Acknowledgements

NR, JE and SOR acknowledge financial support by the Bundesministerium für Bildung und Forschung (BMBF) of Germany via project NG-FLIM (project number 13N15327 and 13N15328). JIG acknowledges financial support from the European Union’s Horizon 2021 research and innovation program under the Marie Sk-lodowskaCurie Grant Agreement no. 101062508 (project name: SOADOPP). JE and JIG acknowledge financial support by the DFG through Germany’s Excellence Strategy EXC 2067/1-390729940. JE and ON thank the European Research Council (ERC) for financial support via project “smMIET” (grant agreement no. 884488) under the European Union’s Horizon 2020 research and innovation program. We thank Alexey Chizhik for providing us with the illustration of our ISM microscope setup, and we thank Anna Chizhik for carefully reading the manuscript.

## Author Contributions Statement

NR and ON built the setup, performed the microscopic imaging, and analyzed the data. JIG prepared the samples, performed the microscopic imaging and analyzed the data. JCT helped with the implementation of the ISM plugin for the analysis software. IG helped with setup and experiments. JE conceived the idea, worked out the theoretical framework, and helped with data analysis. NR, ON, JIG, SOR, and JE wrote the manuscript and did the final editing.

## References

[1] Betzig, E., Patterson, G.H., Sougrat, R., Lindwasser, O.W., Olenych, S., Bonifacino, J.S., Davidson, M.W., Lippincott-Schwartz, J., Hess, H.F.: Imaging intracellular fluorescent proteins at nanometer resolution. Science 313(5793), 1642–1645 (2006) 10.1126/science.1127344

[2] Huang, B., Wang, W., Bates, M., Zhuang, X.: Three-dimensional super-resolution imaging by stochastic optical reconstruction microscopy. Science 319(5864), 810–813 (2008) 10.1126/science.1153529

[3] Heilemann, M., Linde, S., Schüttpelz, M., Kasper, R., Seefeldt, B., Mukherjee, A., Tinnefeld, P., Sauer, M.: Subdiffraction-resolution fluorescence imaging with conventional fluorescent probes. Angewandte Chemie International Edition 47(33), 6172–6176 (2008) 10.1002/anie.200802376

[4] Sharonov, A., Hochstrasser, R.M.: Wide-field subdiffraction imaging by accumulated binding of diffusing probes. Proceedings of the National Academy of Sciences 103(50), 18911–18916 (2006) 10.1073/pnas.06096431

[5] Schnitzbauer, J., Strauss, M.T., Schlichthaerle, T., Schueder, F., Jungmann, R.: Super-resolution microscopy with DNA-PAINT. Nature Protocols 12(6), 1198–1228 (2017) 10.1038/nprot.2017.024

[6] Reinhardt, S.C., Masullo, L.A., Baudrexel, I., Steen, P.R., Kowalewski, R., Eklund, A.S., Strauss, S., Unterauer, E.M., Schlichthaerle, T., Strauss, M.T., et al.: Ångström-resolution fluorescence microscopy. Nature 617(7962), 711–716 (2023) 10.1038/s41586-023-05925-9

[7] Thiele, J.C., Helmerich, D.A., Oleksiievets, N., Tsukanov, R., Butkevich, E., Sauer, M., Nevskyi, O., Enderlein, J.: Confocal fluorescence-lifetime singlemolecule localization microscopy. ACS nano 14(10), 14190–14200 (2020) 10.1021/acsnano.0c07322

[8] Oleksiievets, N., Mathew, C., Thiele, J.C., Gallea, J.I., Nevskyi, O., Gregor, I., Weber, A., Tsukanov, R., Enderlein, J.: Single-molecule fluorescence lifetime imaging using wide-field and confocal-laser scanning microscopy: A comparative analysis. Nano Letters 22(15), 6454–6461 (2022) 10.1021/acs.nanolett.2c01586

[9] Chizhik, A.I., Rother, J., Gregor, I., Janshoff, A., Enderlein, J.: Metal-induced energy transfer for live cell nanoscopy. Nature Photonics 8(2), 124–127 (2014) 10.1038/nphoton.2013.345

[10] Isbaner, S., Karedla, N., Kaminska, I., Ruhlandt, D., Raab, M., Bohlen, J., Chizhik, A., Gregor, I., Tinnefeld, P., Enderlein, J., et al.: Axial colocalization of single molecules with nanometer accuracy using metal-induced energy transfer. Nano Letters 18(4), 2616–2622 (2018) 10.1021/acs.nanolett.8b00425

[11] Thiele, J.C., Jungblut, M., Helmerich, D.A., Tsukanov, R., Chizhik, A., Chizhik, A.I., Schnermann, M.J., Sauer, M., Nevskyi, O., Enderlein, J.: Isotropic threedimensional dual-color super-resolution microscopy with metal-induced energy transfer. Science Advances 8(23), 2506 (2022) 10.1126/sciadv.abo2506

[12] Sheppard, C.R.: Super-resolution in confocal imaging. Optik (Stuttgart) 80(2), 53–54 (1988)

[13] Müller, C.B., Enderlein, J.: Image scanning microscopy. Physical Review Letters 104(19), 198101 (2010) 10.1103/PhysRevLett.104.198101

[14] Sheppard, C.J., Mehta, S.B., Heintzmann, R.: Superresolution by image scanning microscopy using pixel reassignment. Optics Letters 38(15), 2889–2892 (2013) 10.1364/OL.38.002889

[15] Sheppard, C.J., Castello, M., Tortarolo, G., Vicidomini, G., Diaspro, A.: Image formation in image scanning microscopy, including the case of two-photon excitation. Journal of the Optical Society of America A 34(8), 1339–1350 (2017) 10.1364/JOSAA.34.001339

[16] Sheppard, C.J., Castello, M., Tortarolo, G., Deguchi, T., Koho, S.V., Vicidomini, G., Diaspro, A.: Pixel reassignment in image scanning microscopy: a re-evaluation. Journal of the Optical Society of America A 37(1), 154–162 (2020) 10.1364/JOSAA.37.000154

[17] Kner, P., Chhun, B.B., Griffis, E.R., Winoto, L., Gustafsson, M.G.: Superresolution video microscopy of live cells by structured illumination. Nature Methods 6(5), 339–342 (2009) 10.1038/nmeth.1324

[18] Reymond, L., Ziegler, J., Knapp, C., Wang, F.-C., Huser, T., Ruprecht, V., Wieser, S.: SIMPLE: Structured illumination based point localization estimator with enhanced precision. Optics Express 27(17), 24578–24590 (2019) 10.1364/OE.27.024578

[19] Reymond, L., Huser, T., Ruprecht, V., Wieser, S.: Modulation-enhanced localization microscopy. Journal of Physics: Photonics 2(4), 041001 (2020) 10.1088/2515-7647/ab9eac

[20] Cnossen, J., Hinsdale, T., Thorsen, R.Ø., Siemons, M., Schueder, F., Jungmann, R., Smith, C.S., Rieger, B., Stallinga, S.: Localization microscopy at doubled precision with patterned illumination. Nature Methods 17(1), 59–63 (2020) 10.1038/s41592-019-0657-7

[21] Sun, Y., Yin, L., Cai, M., Wu, H., Hao, X., Kuang, C., Liu, X.: Modulated illumination localization microscopy-enabled sub-10 nm resolution. Journal of Innovative Optical Health Sciences 15(02), 2230004 (2022) 10.1142/S179354582230004X

[22] Kalisvaart, D., Cnossen, J., Hung, S.-T., Stallinga, S., Verhaegen, M., Smith, C.S.: Precision in iterative modulation enhanced single-molecule localization microscopy. Biophysical Journal 121(12), 2279–2289 (2022) 10.1016/j.bpj.2022.05.027

[23] York, A.G., Chandris, P., Nogare, D.D., Head, J., Wawrzusin, P., Fischer, R.S., Chitnis, A., Shroff, H.: Instant super-resolution imaging in live cells and embryos via analog image processing. Nature Methods 10(11), 1122–1126 (2013) 10.1038/nmeth.2687

[24] Schulz, O., Pieper, C., Clever, M., Pfaff, J., Ruhlandt, A., Kehlenbach, R.H., Wouters, F.S., Großhans, J., Bunt, G., Enderlein, J.: Resolution doubling in fluorescence microscopy with confocal spinning-disk image scanning microscopy. Proceedings of the National Academy of Sciences 110(52), 21000–21005 (2013) 10.1073/pnas.1315858110

[25] Tenne, R., Rossman, U., Rephael, B., Israel, Y., Krupinski-Ptaszek, A., Lapkiewicz, R., Silberberg, Y., Oron, D.: Super-resolution enhancement by quantum image scanning microscopy. Nature Photonics 13(2), 116–122 (2019) 10.1038/s41566-018-0324-z

[26] Rossman, U., Tenne, R., Solomon, O., Kaplan-Ashiri, I., Dadosh, T., Eldar, Y.C., Oron, D.: Rapid quantum image scanning microscopy by joint sparse reconstruction. Optica 6(10), 1290–1296 (2019) 10.1364/OPTICA.6.001290

[27] Castello, M., Tortarolo, G., Buttafava, M., Deguchi, T., Villa, F., Koho, S., Pesce, L., Oneto, M., Pelicci, S., Lanzanó, L., et al.: A robust and versatile platform for image scanning microscopy enabling super-resolution flim. Nature Methods 16(2), 175–178 (2019) 10.1038/s41592-018-0291-9

[28] Buttafava, M., Villa, F., Castello, M., Tortarolo, G., Conca, E., Sanzaro, M., Piazza, S., Bianchini, P., Diaspro, A., Zappa, F., et al.: SPAD-based asynchronous-readout array detectors for image-scanning microscopy. Optica 7(7), 755–765 (2020) 10.1364/OPTICA.391726

[29] Koho, S.V., Slenders, E., Tortarolo, G., Castello, M., Buttafava, M., Villa, F., Tcarenkova, E., Ameloot, M., Bianchini, P., Sheppard, C.J., et al.: Two-photon image-scanning microscopy with SPAD array and blind image reconstruction. Biomedical Optics Express 11(6), 2905–2924 (2020) 10.1364/BOE.374398

[30] Slenders, E., Perego, E., Buttafava, M., Tortarolo, G., Conca, E., Zappone, S., Pierzynska-Mach, A., Villa, F., Petrini, E.M., Barberis, A., et al.: Cooled SPAD array detector for low light-dose fluorescence laser scanning microscopy. Biophysical Reports 1(2), 100025 (2021) 10.1016/j.bpr.2021.100025

[31] Roth, S., Sheppard, C.J., Wicker, K., Heintzmann, R.: Optical photon reassignment microscopy (OPRA). Optical Nanoscopy 2(1), 1–6 (2013) 10.1186/2192-2853-2-5

[32] Roth, S., Sheppard, C.J., Heintzmann, R.: Superconcentration of light: circumventing the classical limit to achievable irradiance. Optics Letters 41(9), 2109–2112 (2016) 10.1364/OL.41.002109

[33] Fazel, M., Grussmayer, K.S., Ferdman, B., Radenovic, A., Shechtman, Y., Enderlein, J., Pressé, S.: Fluorescence microscopy: a statistics-optics perspective. arXiv (2023) 10.48550/arXiv.2304.01456

[34] Stallinga, S., Radmacher, N., Delon, A., Enderlein, J.: Optimal transfer functions for bandwidth-limited imaging. Physical Review Research 4(2), 023003 (2022) 10.1103/PhysRevResearch.4.023003

[35] Smith, C.S., Slotman, J.A., Schermelleh, L., Chakrova, N., Hari, S., Vos, Y., Hagen, C.W., Müller, M., Cappellen, W., Houtsmuller, A.B., et al.: Structured illumination microscopy with noise-controlled image reconstructions. Nature Methods 18(7), 821–828 (2021) 10.1038/s41592-021-01167-7

[36] Holden, S.J., Uphoff, S., Kapanidis, A.N.: Daostorm: an algorithm for highdensity super-resolution microscopy. Nature Methods 8(4), 279–280 (2011) 10.1038/nmeth0411-279

[37] Ovesnỳ, M., Křížek, P., Borkovec, J., Švindrych, Z., Hagen, G.M.: Thunderstorm: a comprehensive imagej plug-in for palm and storm data analysis and superresolution imaging. Bioinformatics 30(16), 2389–2390 (2014) 10.1093/bioinformatics/btu202

[38] Stein, S.C., Thiart, J.: TrackNTrace: A simple and extendable open-source framework for developing single-molecule localization and tracking algorithms. Scientific Reports 6(1), 37947 (2016) 10.1038/srep37947

[39] Linde, S.: Single-molecule localization microscopy analysis with imagej. Journal of Physics D: Applied Physics 52(20), 203002 (2019) 10.1088/1361-6463/ab092f

[40] Stein, S., Thiart, J., Thiele, J.C., Radmacher, N.: TrackNTrace Freeware Package. https://gitlab.gwdg.de/igregor/single-molecule-localization-microscopy-with-image-scanning-microscopy.git (2023)

[41] Rieger, B., Stallinga, S.: The lateral and axial localization uncertainty in superresolution light microscopy. ChemPhysChem 15(4), 664–670 (2014) 10.1002/cphc.201300711

[42] Zwettler, F., Reinhard, S., Gambarotto, D., Bell, T., Hamel, V., Guichard, P., Sauer, M.: Molecular resolution imaging by post-labeling expansion singlemolecule localization microscopy (ex-smlm). Nature Communications 11, 3388 (2020) 10.1038/s41467-020-17086-8

[43] Thiele, J.C., Nevskyi, O., Helmerich, D.A., Sauer, M., Enderlein, J.: Advanced data analysis for fluorescence-lifetime single-molecule localization microscopy. Frontiers in Bioinformatics 1, 740281 (2021) 10.3389/fbinf.2021.740281

[44] Wegel, E., Göhler, A., Lagerholm, B.C., Wainman, A., Uphoff, S., Kaufmann, R., Dobbie, I.: Imaging cellular structures in super-resolution with sim, sted and localisation microscopy: A practical comparison. Scientific Reports 6, 27290 (2016) 10.1038/srep27290

[45] Oleksiievets, N., Sargsyan, Y., Thiele, J.C., Mougios, N., Sograte-Idrissi, S., Nevskyi, O., Gregor, I., Opazo, F., Thoms, S., Enderlein, J., et al.: Fluorescence lifetime DNA-PAINT for multiplexed super-resolution imaging of cells. Communications Biology 5(1), 38 (2022) 10.1038/s42003-021-02976-4

[46] Kalisvaart, D., Hung, S.-T., Smith, C.S.: Quantifying the minimum localization uncertainty of image scanning localization microscopy. Biophysical Reports 4(1) (2024) 10.1016/j.bpr.2024.100143

[47] Dani, A., Huang, B., Bergan, J., Dulac, C., Zhuang, X.: Superresolution imaging of chemical synapses in the brain. Neuron 68(5), 843–856 (2010) 10.1016/j.neuron.2010.11.021

[48] Truckenbrodt, S., Maidorn, M., Crzan, D., Wildhagen, H., Kabatas, S., Rizzoli, S.O.: X10 expansion microscopy enables 25-nm resolution on conventional microscopes. EMBO Reports 19(9), 45836 (2018) 10.15252/embr.201845836

[49] Tortarolo, G., Zunino, A., Fersini, F., Castello, M., Piazza, S., Sheppard, C.J., Bianchini, P., Diaspro, A., Koho, S., Vicidomini, G.: Focus image scanning microscopy for sharp and gentle super-resolved microscopy. Nature Communications 13(1), 7723 (2022) 10.1038/s41467-02235333-y

[50] Slenders, E., Vicidomini, G.: ISM-FLUX: MINFLUX with an array detector. Physical Review Research 5(2), 023033 (2023) 10.1103/PhysRevResearch.5.023033

