## Supporting Information for "Doubling the resolution of fluorescence-lifetime single-molecule localization microscopy with image scanning microscopy"

### Table of contents

1 Supplementary figure: Microtubule cross-sections 2

2 Supplementary figure: iSMLM performance simulation 3

3 Supplementary figure: FL-iMSLM multiplexing 4

4 Supplementary figure: Synaptic cleft 5

### 1 Supplementary figure: Microtubule cross-sections

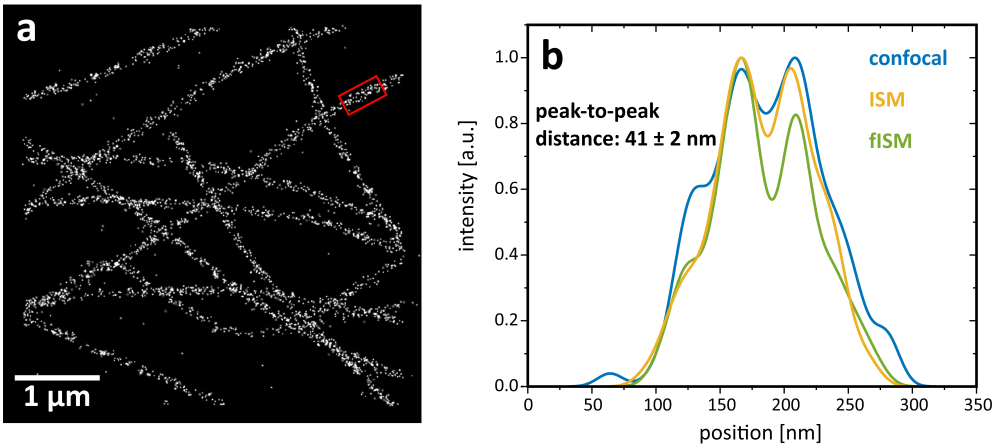


**Fig. S1** Microtubule cross-sections. (a) fISM-dSTORM reconstruction of zoom-in I depicted in Fig. 3(a). (b) Average intensity profile of a microtubule segment shown in (a) for confocal dSTORM, ISM-dSTORM and fISM-dSTORM reconstructions. The segment length is 500 nm and presented in red.

#### 2 Supplementary figure: iSMLM performance simulation

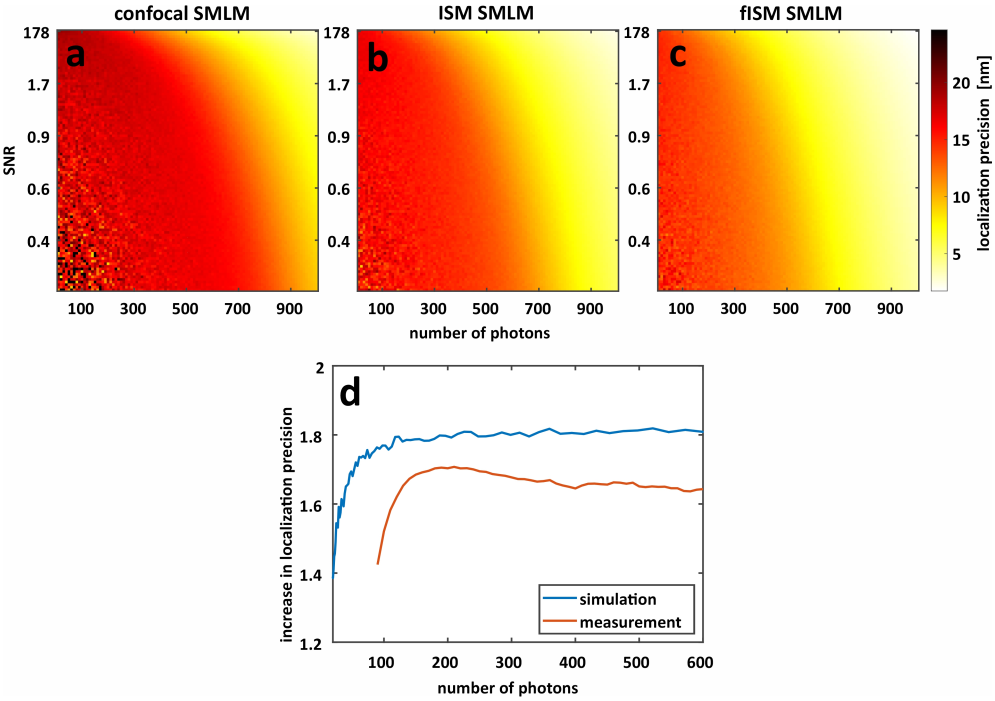


**Fig. S2** iSMLM performance simulation. For a wide range of photon numbers and background noise. Localization precision was determined by SMLM analysis package TrackNTrace. (a) with no ISM pixel reassignment, (b) with pixel reassignment and (c) with addition Fourier reweighting. (d) increase in localization precision between confocal SMLM and iSMLM as a function of minimum number of photons per localizaion.

### 3 Supplementary figure: FL-iMSLM multiplexing

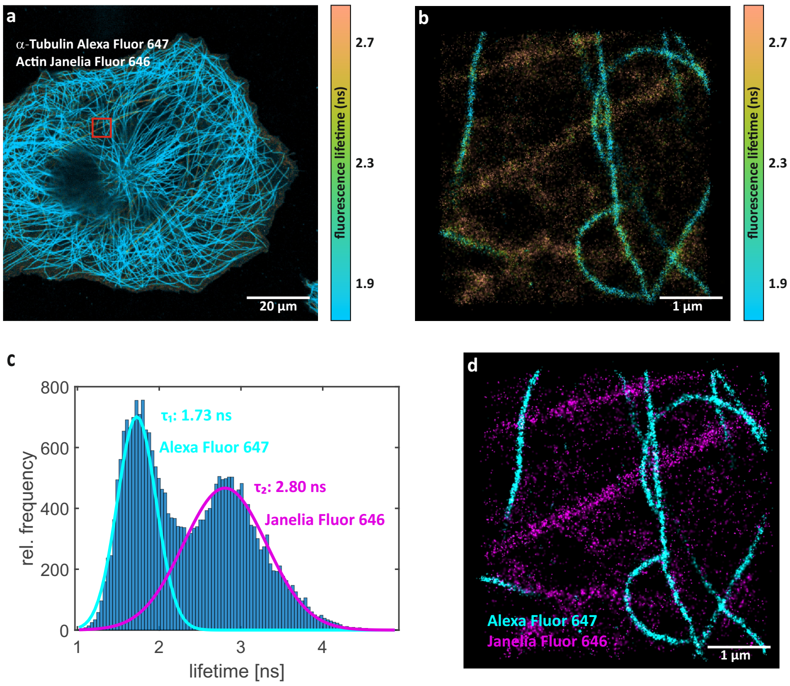


**Fig. S3** dSTORM based FL-iSMLM multiplexing. (a) FL-ISM overview of a COS-7 cell where $\alpha$-tubulin was labeled with Alexa 647 and actin with phalloidin-Janelia Fluor 646 fluorophores. Zoom is the actual 5 µm × 5 µm region of interest of the dSTORM measurement. Colors represent fluorescence lifetime, determined from TCSPC tail fitting. (b) Super-resolved Fourier-reweighted ISM dSTORM reconstruction FLIM image made from $6.5 \cdot 10^4$ images with 10 frame binning. (c) Fitted lifetime histogram based on individual single-molecule localizations for the corresponding region of interest. (d) Dual-color super-resolved image based on the lifetime data. The two lifetime components were separated using lifetime pattern matching classification. Here localizations events are colored cyan if their decay pattern belongs to that of Alexa 647 with a probability of more than 99%. And magenta, if the same is true for the Janelia Fluor 646 pattern.

### 4 Supplementary figure: Synaptic cleft

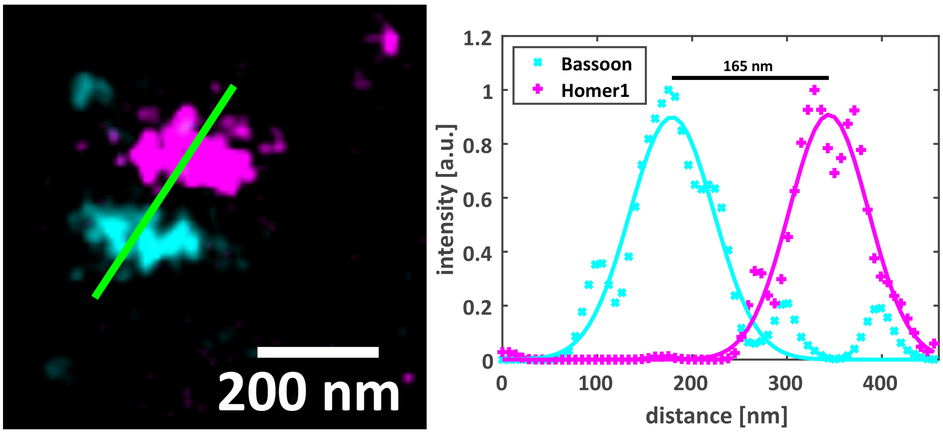


**Fig. S4** An example of one synapse with the distance between Bassoon and Homer1 measured across the synaptic cleft. (a) DNA-PAINT iSMLM reconstruction of zoom-in I depicted in Fig. 5(a). Homer1 was labeled with Atto 565 and is depicted in magenta, Bassoon was stained with Cy3b and is shown in cyan (b) Intensity profile along the line shown in green with individual Gaussian fits for the two proteins. Distance was calculated from the absolute difference between the mean values of the respective Gaussian fits.
